# Linking Molecular Tension and Cellular Tractions: A Multiscale Approach to Focal Adhesion Mechanics

**DOI:** 10.1101/2025.01.09.632081

**Authors:** Samet Aytekin, Laurens Kimps, Quinten Coucke, Débora Linhares, Swaraj Deodhar, Ruth Cardinaels, Mar Cóndor, Jorge Barrasa-Fano, Hans Van Oosterwyck, Susana Rocha

## Abstract

Focal adhesions (FAs) are mechanosensitive structures that mediate force transmission between cells and the extracellular matrix. While Traction Force Microscopy (TFM) quantifies cellular tractions exerted on deformable substrates, Förster Resonance Energy Transfer (FRET)-based tension probes, such as Vinculin Tension Sensors (VinTS), measure molecular-scale forces within FA proteins. Despite their potential synergy, these methods have rarely been combined to explore the interplay between molecular tension and cellular tractions. Here, we introduce a framework integrating TFM and VinTS to investigate FA mechanics across scales. At cell level, tractions and vinculin tension increased with substrate stiffness. At FA level, vinculin tension correlated with vinculin density, while tractions scaled with FA area, total vinculin content and vinculin density. Direct comparison of tractions to tension revealed a complex, heterogenous relationship between these forces, possibly linked to diverse cell and FA maturation states. Sub-FA analysis revealed conserved spatial patterns, with tension and traction increasing towards the cell periphery. This multiscale approach provides insights into the multiscale dynamics of FA mechanotransduction, bridging the gap between molecular forces and cellular mechanics.j

## Introduction

Mechanobiology is an interdisciplinary field that investigates the reciprocal relationship between mechanical forces and biological processes. Mechanotransduction – the process by which cells sense and respond to mechanical signals from their environment – plays a crucial role in fundamental cellular functions such as proliferation, differentiation and migration, as well as in pathological conditions like fibrosis, cancer and cardiovascular diseases^1–5^. Deeper understanding of mechanotransduction offers insights into cellular behavior and disease mechanisms, leading to innovative therapies that restore the mechanical equilibrium in the cellular microenvironment.

Central to mechanotransduction are focal adhesions (FAs), multifunctional complexes at the plasma-membrane that physically connect the actin cytoskeleton to the extracellular matrix (ECM). These structures are pivotal for sensing mechanical cues and translating them into cellular responses. Through actomyosin-mediated contractility, FAs transmit forces from the cytoskeleton to the ECM, allowing cells to sense their surroundings and convert biophysical cues into cellular processes such as cell migration, differentiation, cell-cycle progression and cell death^6,7^. FAs exhibit a complex structure and dynamic protein composition, undergoing changes in both morphology and composition based on their maturation stage^8,9^. Vinculin, a mechanosensitive protein, is essential for FA maturation and strengthening. It transitions from an auto-inhibited state to an active form through ligand-assisted unfolding and stretching^10–13^. Once activated, vinculin interacts with other FA proteins such as actin, paxillin and talin to mediate mechanosensing and lamellipodial actin dynamics^14–17^. These roles make vinculin is a key player in cell adhesion, migration, and mechanotransduction^18,19^.

One of the most widely used methods to quantify forces transmitted by FAs at the cell-ECM interface is Traction Force Microscopy (TFM). Traditionally applied to 2D *in vitro* cultures, TFM involves seeding the cells on a synthetic, ECM-like substrate^20–22^. The substrate, typically an elastic hydrogel with embedded fiducial markers, is imaged using optical microscopy to capture two states: a stressed state, where cells are mechanically active, and stress-free or relaxed state, where cells have either been removed or actomyosin contractility has been disrupted. Image processing algorithms track the movement of fiducial markers between these states to measure hydrogel deformation. Using the known mechanical properties of the hydrogel, mechanical models can infer cell-generated forces and tractions (force per unit of area) from these displacements. Unlike cantilever or micropillar-based TFM^23,24^, hydrogel-based TFM provides a continuous substrate for cellular adhesion, allowing cells to developed FA morphologies that are not restricted by a certain cross-sectional area. This approach also allows for the computation of discretized traction fields, theoretically limited only by the optical resolution of the microscope.

While TFM is instrumental in quantifying and visualizing the tractions transmitted via FAs, it does not provide a direct readout of the molecular tension within FA proteins, as it captures tractions exerted at the substrate level. Förster Resonance Energy Transfer (FRET)-based molecular tension probes address this limitation by directly measuring forces at the molecular scale^25^. These probes detect conformational changes by measuring the changes in FRET – a phenomenon where energy transfers non-radiatively between two nearby fluorophores – in response to mechanical force^26^. FRET-based tension probes consist of a pair of fluorescent proteins (*Donor* and *Acceptor*) linked by a short, flexible polypeptide that undergoes conformational changes upon tensional forces. When no tension is applied, the fluorescent proteins remain in close proximity and show high-FRET efficiency. When the linker is stretched under tension, the distance between the fluorescent proteins increases, leading to a decrease in FRET. This force-induced separation of the FRET-pair and subsequent change in FRET allows for sensitive detection of molecular forces in the pN range, making FRET-based tension probes powerful for studying mechanobiology at the molecular scale. Among the FRET-based tension probes, Vinculin Tension Sensor (VinTS) is the best characterized and has been extensively used to investigate FA mechanics^9,24,25,27–33^. By incorporating the FRET tension probes within vinculin, VinTS reveals the magnitude and spatial distribution of forces within FAs, offering a molecular view on force transmission at cell-matrix interfaces.

While TFM and FRET-based tension probes are powerful techniques on their own, they offer insights into cell-matrix forces at different levels. TFM quantifies relatively large forces that are sufficient to induce measurable substrate deformations, depending on factors like substrate stiffness. These forces originate from subcellular regions, such as FAs and lamellipodia. In contrast, FRET-based tension probes report the molecular forces at the nanoscale. To date, only two studies have combined FRET tension probes with cell-ECM force measurements. Chang et al. (2013) performed TFM on cells expressing VinTS and reported a slight inverse linear correlation between mean traction and mean FRET ratio at FAs. However, their analysis was limited by the low-throughput of just four cells^27^. Sarangi et al. (2017) simultaneously mapped traction forces using a micropillar array and vinculin tension within FAs. This approach revealed how changes in traction forces corresponded to variations in vinculin tension^24^. However, their *in vitro* system confined FAs to the top of micropillars, which may not accurately reflect native FA behavior and could influence their dynamics or functionality.

In this study, we established an experimental and computational framework integrating hydrogel-based TFM, which enables FA formation on an unrestrictive, continuous substrate, with VinTS to investigate the interplay between vinculin tension and traction forces at FAs. Our findings showed that tractions increase with FA area, vinculin content and density, while vinculin tension was mainly regulated by vinculin density. At cell-level, we observed both tension and traction magnitude increase with increasing substrate stiffness. At FA-level, we identified three correlation patterns between vinculin tension and traction – direct, inverse and uncorrelated – suggesting varied cellular and FA states. At sub-FA level, vinculin tension and traction followed a similar trend along the longitudinal axis of FAs, consistent across different cell and FA populations. The framework advances our understanding of subcellular mechanotransduction and provides a robust foundation for future studies of cellular force dynamics and mechanobiology.

## Results

### Integrated Workflow for TFM and FRET-based tension probes

To integrate TFM with FRET-based tension probes, we developed a custom protocol to overcome the technical challenges of combining these methods (Fig. 1a, details in the methods section). Sub-cellular FRET imaging requires high numerical aperture (NA) objectives, which have a limited working distance of ~300 μm. To ensure that cells on the polyacrylamide (PAA) gel remained within the objective’s working distance, we fabricated thin, defect-free PAA substrates using 250 μm thick silicone spacers and #0 coverslips. The spacers also created flat gel surfaces, aligning all focal adhesions within the same z-plane, thereby facilitating FRET imaging. To calculate tractions exerted by individual mouse embryonic fibroblasts (MEFs) accurately, we maintained a low cell density of ~25 cells/mm^2^. At this density, cell spreading required optimal collagen coating, which was achieved by using an acidic solution^34^. VinTS-transfected vinculin double knock out MEFs are seeded on the collagen-coated gels and allowed to spread overnight before imaging.

**Figure 1.**
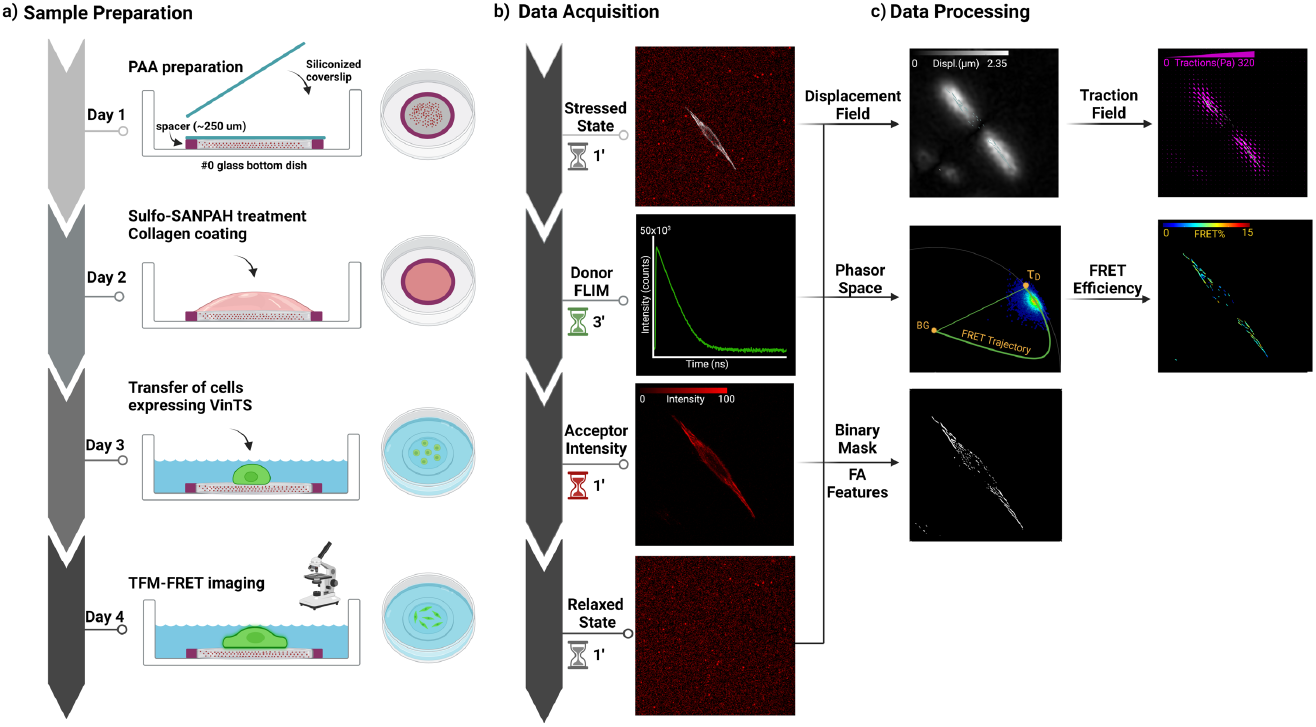
Experimental workflow and analysis pipeline for investigating FA forces. (**a**) Schematic overview of the experimental procedure: Day 1 – preparation of thin (250 μm) and flat PAA gels; Day 2 – surface functionalization with Sulfo-SANPAH and collagen coating in acidic solution; Day 3 –seeding of cells transiently transfected with VinTS plasmid; Day 4 – live cell imaging. (**b**) Imaging and data acquisition on day 4: stressed state bead and cell image acquisition (TFM imaging) followed by FLIM data acquisition of donor (*Clover*) channel (FRET imaging) and intensity data of acceptor (*mRuby2*) channel. Relaxed state bead image acquired after the removal of the cells and the relaxation of the gel using a mild SDS solution. (**c**) Data processing: displacement fields generated from the overlay images of stressed and relaxed state bead images, and subsequently converted into tractions. FRET efficiency is derived from the FRET trajectory line on the phasor space. FA structural and molecular properties are quantified from the acceptor intensity data after applying a binary FA mask. *Illustrations were created using BioRender*.*com*.

Similar to sample preparation, combining TFM and FRET-probes required a specific data acquisition protocol (Fig. 1b). Due to the dynamic behavior of fibroblasts, which induces relatively fast changes in cell morphology, it is crucial to minimize the time interval between the TFM and FRET data acquisition. First, bead images of the stressed state were acquired, followed by Fluorescence Lifetime Imaging Microscopy (FLIM) images of the donor channel (used to calculate FRET efficiency)^35–38^. To analyze the vinculin content and the morphology of the FAs, fluorescence image of the acceptor channel was then acquired. These steps were repeated for ~5-10 cells in the sample. Next, cells were removed using sodium dodecyl sulfate (SDS), allowing the relaxation of the PAA gel, and bead images of the relaxed state were acquired for all measured cells, completing the data acquisition process.

Stressed and relaxed state bead images were subsequently analyzed using a previously established 2D TFM workflow^39^. Displacement fields were calculated using Free Form Deformation (FFD) image registration of bead positions between the two states^40^ (Fig. 1c, top row). Traction fields were derived from the displacement fields using a Tikhonov-regularized Fourier transform traction cytometry algorithm (Fig.1c, top row)^21,39^. To calculate the FRET efficiency of the vinculin tension probes, fluorescence lifetimes were analyzed using the phasor approach^41–45^. Phasor-transformation of the donor-specific FLIM data can be used to directly retrieve FRET efficiencies from a so-called FRET trajectory line. This line extends from the average phasor position of the donor in the absence of acceptor (0% FRET), calculated from Clover-only transfected cells, to the position of background signal (100% FRET), obtained from non-transfected cells (Fig. 1c, middle row). The FRET efficiency of VinTS-expressing cells was then calculated from the position of the acquired photons on the FRET trajectory line. Since the calibration of the VinTS, which converts FRET efficiency into tensile force, was done *in silico*^29^, we chose to report FRET efficiencies without converting them to absolute force values. TFM, FRET efficiencies and acceptor intensity data were integrated using a custom MATLAB code. A binary mask created by Ilastik^46^ (Fig. 1c, bottom row) was used to segment the FAs, allowing each FA to be assigned a traction, FRET and intensity information, together with morphological features such as FA area and orientation.

### Cell tractions and vinculin tension increase with stiffness at cell level

To evaluate how substrate stiffness influences cellular forces, we analyzed cell-averaged values of tractions, FRET efficiencies, and FA properties in MEFs seeded on PAA gels (Fig. 2a). Experiments were performed on gels with a Young’s modulus of 4.35 ± (s.d.) 0.20 kPa (referred to as 4.5 kPa) and 13.10 ± 0.23 kPa (referred to as 13 kPa), as characterized by shear rheological measurements. These stiffness values were chosen to ensure the substrates were stiff enough to support cell spreading but soft enough to undergo detectable deformations. Cells on the stiffer substrate spread more (Fig. 2b) and exerted higher tractions (Fig. 2c), indicating enhanced cell-ECM interactions and cell contractility with increasing stiffness. To investigate whether increased tractions correlated with changes in molecular tension within FAs, we determined cell-averaged FRET efficiencies of FAs in VinTS-expressing cells on soft and stiff PAA gels, as well as on glass. FRET efficiency decreased with increasing stiffness: highest on the soft gel, intermediate on the stiff gel, and lowest on glass (Fig. 2f). To validate these results, we used a cytosolic, force-insensitive probe, Tension Sensing Module (TSMod, Fig. 2d), which exhibited high FRET efficiencies across all conditions (Fig. 2f). These results confirmed that the observed change in VinTS FRET efficiency was due to the change of tension across vinculin. The progressive FRET decrease indicates that vinculin tension increases with stiffness, suggesting a possible link between traction and FA tension.

**Figure 2.**
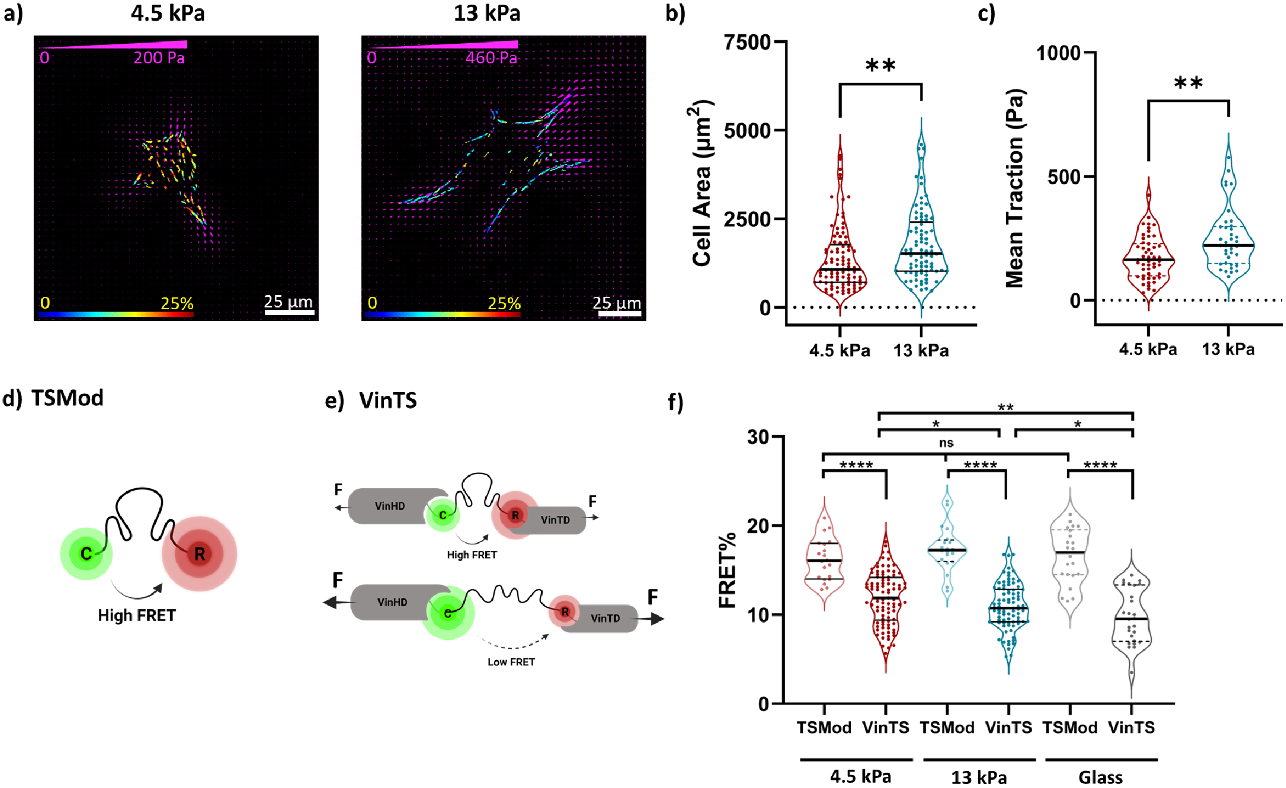
Effect of substrate stiffness on the cell size, cellular tractions and vinculin tension. (**a**) Traction and FRET overlay images of representative cells on soft (4.5 kPa) and stiff (13 kPa) PAA substrate. Traction arrows are scaled with traction magnitude. (**b**) Comparison of cell surface area between soft and stiff PAA substrate. n = 93 and 85 cells. (**c**) Comparison of cell-averaged tractions between the soft and stiff substrate. n = 51 and 38 cells. (**d**) Illustration of the structure and working principle of Tension Sensing Module (TSMod) during donor excitation; Clover (C) and mRuby2 (R) FRET pair is separated with a synthetic polypeptide (GGSGGS)_7_; FRET efficiency remains high due to inability to support tension. (**e**) Illustration of the structure and working principle of Vinculin Tension Sensor (VinTS); TSMod is inserted between vinculin head domain (VinHD) and vinculin tail domain (VinTD); FRET efficiency decreases as a result of applied tension. (**f**) Averaged FA FRET efficiencies per cell for VinTS and force-insensitive control TSMod compared on three different substrates. n = 20, 93, 29, 85, 22 and 26 cells. Statistical analysis between two groups were performed using nonparametric Mann-Whitney U-test for (b) and (c), and unpaired two-tailed parametric Student’s t-test for (f): *p < 0.05, **p < 0.01, ****p < 0.0001. *Illustrations (d) and (e) were created using BioRender*.*com*.

Since FAs are the primary structures where cellular forces are transmitted, we investigated whether changes in cell tractions and vinculin tension are associated with FA structural (number, total area, density, size, eccentricity and orientation) and molecular (total vinculin content and vinculin density) properties (Supplementary Fig. 1). Interestingly, we found none of the tested properties significantly altered between these two selected stiffnesses.

### Tractions and vinculin tension depend on FA structural and molecular properties

Since FA structural and molecular properties were consistent across different stiffnesses, only data from 4.5 kPa gels are presented. To investigate whether the FA forces are locally linked to structural and molecular properties, we evaluated traction and vinculin tension at individual FA level.

The mean traction values of each FA (Fig. 3a) were analyzed in relation to FA area, orientation, total vinculin content and vinculin density (Fig. 3d-g). Small FAs and those with lower total vinculin content or density exhibited a wide range of traction values, while larger FAs and those with higher vinculin content or density were consistently associated with higher traction values. Notably, an upper limit for traction was observed, beyond which further increases in FA area, vinculin content or density did not seem to affect traction magnitude. We also observed that tractions tend to be larger for radially oriented FAs (i.e. FAs pointing towards the cell centroid, Fig. 3e). However, this data displayed substantial variability across the entire orientation range, suggesting that orientation with respect to the centroid of the cell is a poor predictor of tractions.

**Figure 3.**
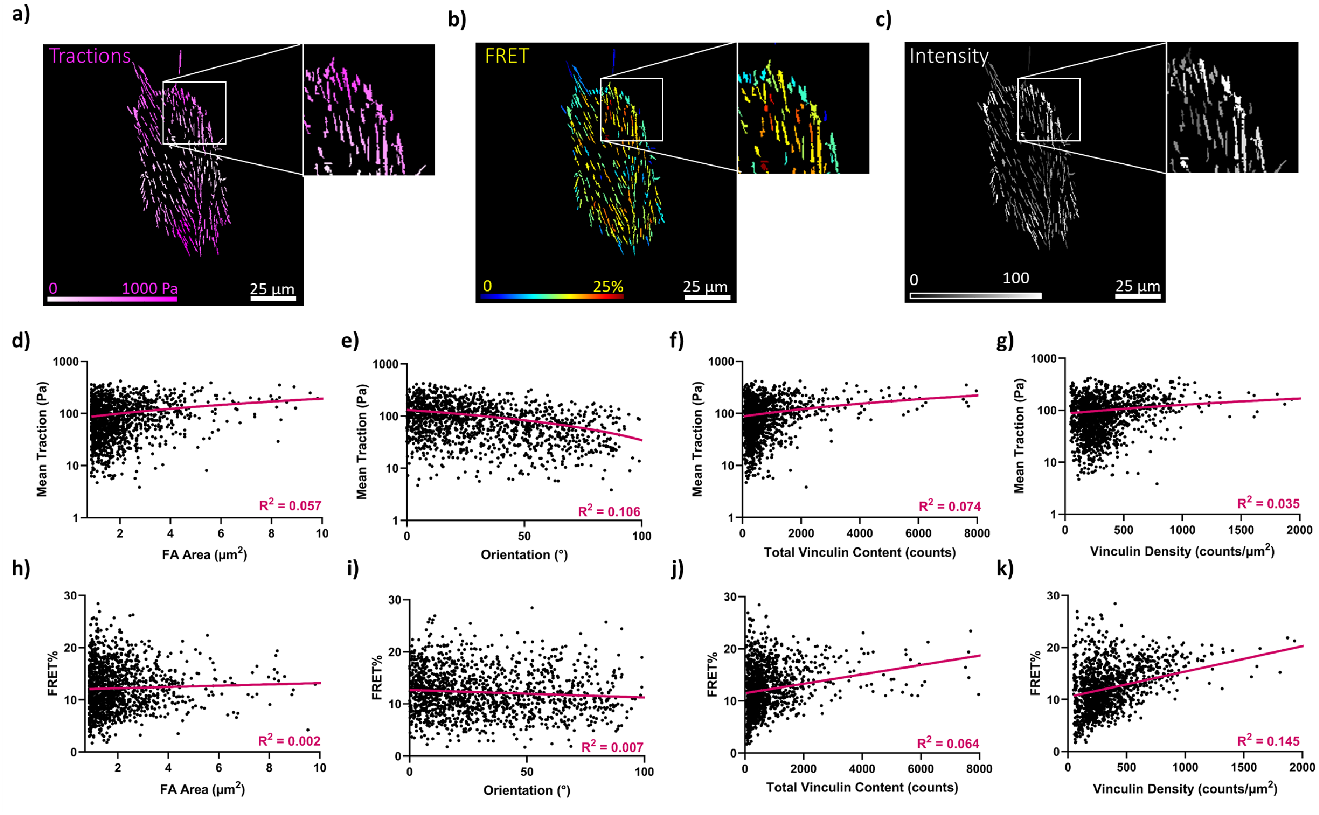
Effect of primary FA structural and molecular properties on mean tractions and FRET efficiency. A mean value of traction (**a**), FRET efficiency (**b**) and intensity (**c**) was assigned to each segmented FA. FA area, orientation, total vinculin content and vinculin density were compared to mean tractions (**d-g**) and mean FRET efficiency (**h-k**). n = 1359 FAs from 28 cells seeded on soft (4.5 kPa) substrate. Purple lines represent the linear regression fit, with R^2^ values indicated on each graph.

To analyze vinculin tension at individual FA level, mean FRET efficiencies were assigned to individual FAs (Fig. 3b) and compared with the same FA properties (Fig. 3h-k). Similar to traction results, we observed that small FAs are associated with a wide range of FRET efficiencies, from very low to very high, while larger FAs are associated with moderate FRET efficiencies, indicating that vinculin tension stabilizes as FAs mature. While a similar distribution was observed for FAs with lower total vinculin content or vinculin density, we observed an increase in FRET efficiency with vinculin density and, to a lesser extent, total vinculin content. The former suggests that the tension over individual vinculin molecules decreases as the local vinculin density increases. No correlation was found between FRET efficiencies and FA orientation (Fig. 3i), indicating the tension over the vinculin does not depend on the direction of the contractile forces. Overall, these results suggest that FA tractions are mainly regulated by FA area, total vinculin content and density, while vinculin tension is mainly regulated by local vinculin density, and to a lesser extent by total vinculin content.

### Correlating vinculin tension to traction at FA level reveals cell-to-cell heterogeneity

At cell level, both cell-averaged tractions and vinculin tension increase with substrate stiffness (Figs. 2c and f), suggesting a positive correlation between traction and vinculin tension. However, neither showed a clear correlation with FA morphology. We next examined whether tractions and vinculin tension are correlated at individual FA level (Fig. 4).

**Figure 4.**
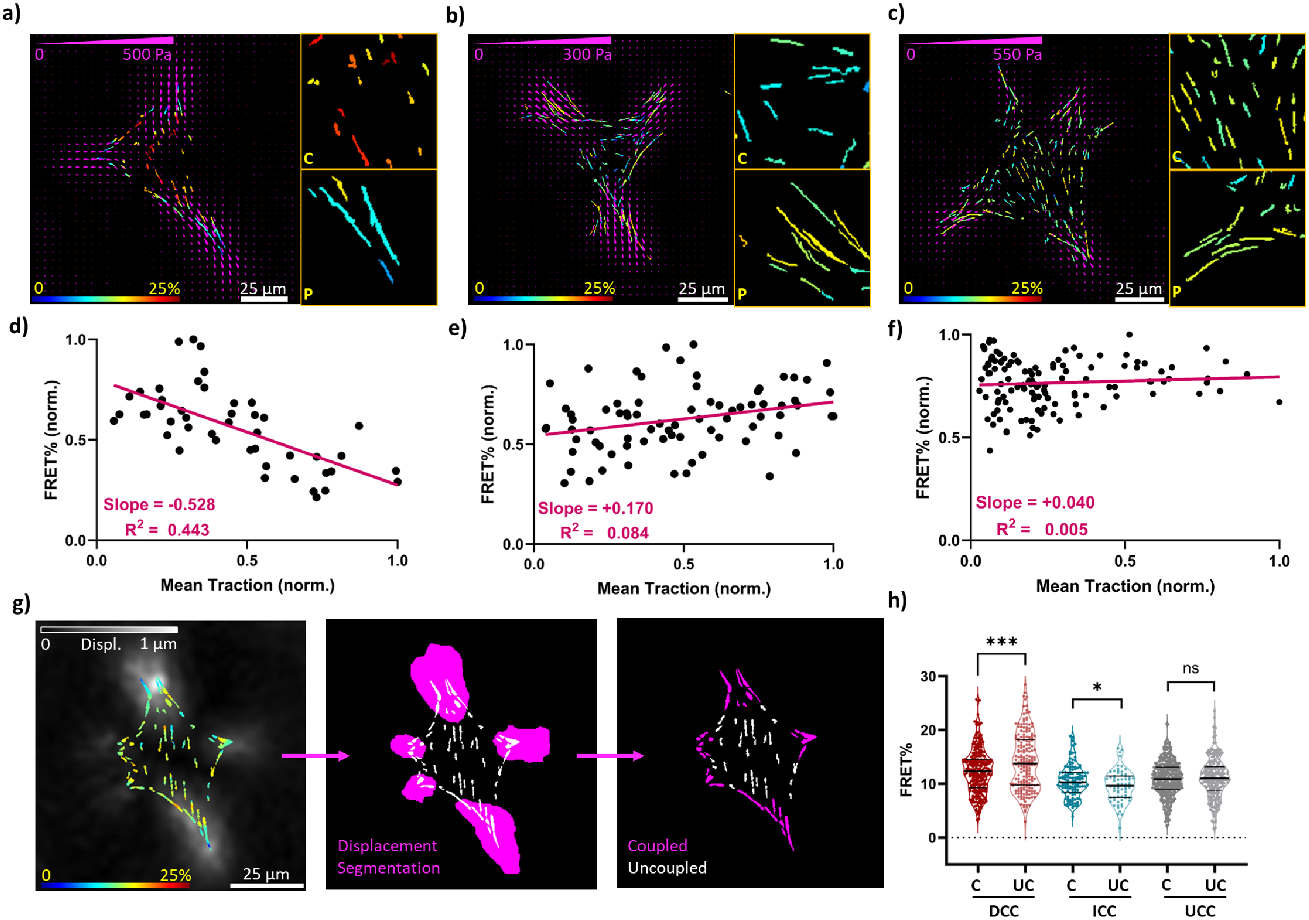
Correlative assessment of FA tractions and vinculin tension. Traction/FRET overlay images of representative cells for each type of correlation: (**a**) directly correlated cell (DCC), (**b**) inversely correlated cell (ICC) and (**c**) uncorrelated cell (UCC); insets with enlarged images show the central (C) and peripheral (P) FAs. (**d-f**) Corresponding FRET efficiency vs mean traction scatter plots of the representative cells shown in (a-c). Purple lines represent the linear regression fit, with slope and R^2^ values indicated on the graphs. (**g**) Displacement-based region selection to segregate the FAs depending on their engagement in the traction generation. (**h**) FRET efficiency distribution between the coupled and uncoupled FAs compared for DCC, ICC and UCC on soft (4.5 kPa) substrate. n = 20, 9 and 22 cells for DCC, ICC and UC, respectively. Statistical analysis between two groups were performed using unpaired two-tailed parametric Student’s t-tests: *p < 0.05 and ***p < 0.001.

Similar to the previous section, tractions and FRET efficiency were averaged for each FA. While no correlation was observed when the data from all cells was pooled (Supplementary Fig. 2a), cell-specific analysis revealed distinct traction-vinculin tension correlations across individual cells (representative cells in Fig. 4a-c and corresponding correlations Fig. 4d-f). To classify these correlations, cells were grouped as directly correlated (negative slope smaller than −0.1 in FRET versus traction graphs; noting that an inverse correlation with FRET efficiency indicates a direct correlation with vinculin tension), inversely correlated (positive slope larger than 0.1), or uncorrelated (slope between −0.1 and 0.1; Supplementary Fig. 2b). Tractions and FRET values were normalized to their maximum *per cell*, confining slope values within −1 to 1. Among the cells analyzed on the soft gel, 39% exhibited a direct correlation, where higher traction coincided with higher vinculin tension (and therefore lower FRET efficiency, Supplementary Fig. 2c). Only 18% of the cells displayed an inverse correlation, where larger tractions corresponded to lower vinculin tension across individual FAs. Notably, 43% of the cells showed no correlation between mean traction and vinculin tension. To understand the effect of stiffness on the different correlation types, the analysis was repeated for cells seeded on the stiff (13 kPa) substrate. We observed an increase in both directly correlated cells (46%) and uncorrelated cells (51%), and a sharp decrease in indirectly correlated cells, with a rare occurrence of of 2%, indicating that the correlation between traction and vinculin tension at individual FA level is indeed affected by substrate stiffness (Supplementary Fig. 2d).

We investigated whether different correlation types were linked to the traction magnitude. However, comparing cell-averaged tractions between directly and inversely correlated cells did not show a significant difference (Supplementary Fig. 2e). Upon close examination of individual FRET images, we observed distinct patterns in FRET efficiencies between the central and peripheral FAs. In cells with a direct vinculin tension-traction relationship, peripheral FAs showed markedly lower FRET than central FAs, suggesting higher vinculin tension at the cell periphery (Fig. 4a). Conversely, in cells with inverse vinculin tension-traction relationship, FRET appeared elevated in peripheral FAs relative to the central ones, suggesting lower vinculin tension at the cell periphery (Fig. 4b). Cells without a clear vinculin tension-traction relationship showed no noticeable differences in FRET between central and peripheral FAs (Fig. 4c). We hypothesized that, in cells with direct correlation, vinculin tension is elevated in FAs actively participating in traction generation. In contrast, cells showing inverse correlation may experience higher vinculin tension in FAs not directly engaged in traction. To test this hypothesis, we overlaid FRET and TFM displacement images, and categorized FAs as “coupled” (associated with areas of elevated displacements) or “uncoupled” (outside the areas with elevated displacements; Fig. 4g). Comparison of FRET efficiencies between these two subsets showed significantly higher vinculin tension (lower FRET efficiency) in coupled FAs in directly correlated cells, while inversely correlated cells exhibited the opposite trend; the vinculin tension was higher in the uncoupled FAs. Cells without a clear FRET-traction relationship showed no significant differences between coupled and uncoupled FAs (Fig. 4h).

### Traction and vinculin tension exhibit similar spatial distributions along individual FAs

Previous studies have shown a spatial distribution of vinculin tension along FAs^9,24,29,30^. To explore how these changes relate to tractions and vinculin density, we analyzed all three variables at the sub-FA level. To account for variability between FAs and cells, each variable was normalized to its maximum and minimum value within each FA (Supplementary Fig. 3). The cell-averaged spatial distributions of intensity, FRET efficiency and tractions were then compared across directly correlated, inversely correlated and uncorrelated cells (as defined in the previous section; Fig. 5a).

**Figure 5.**
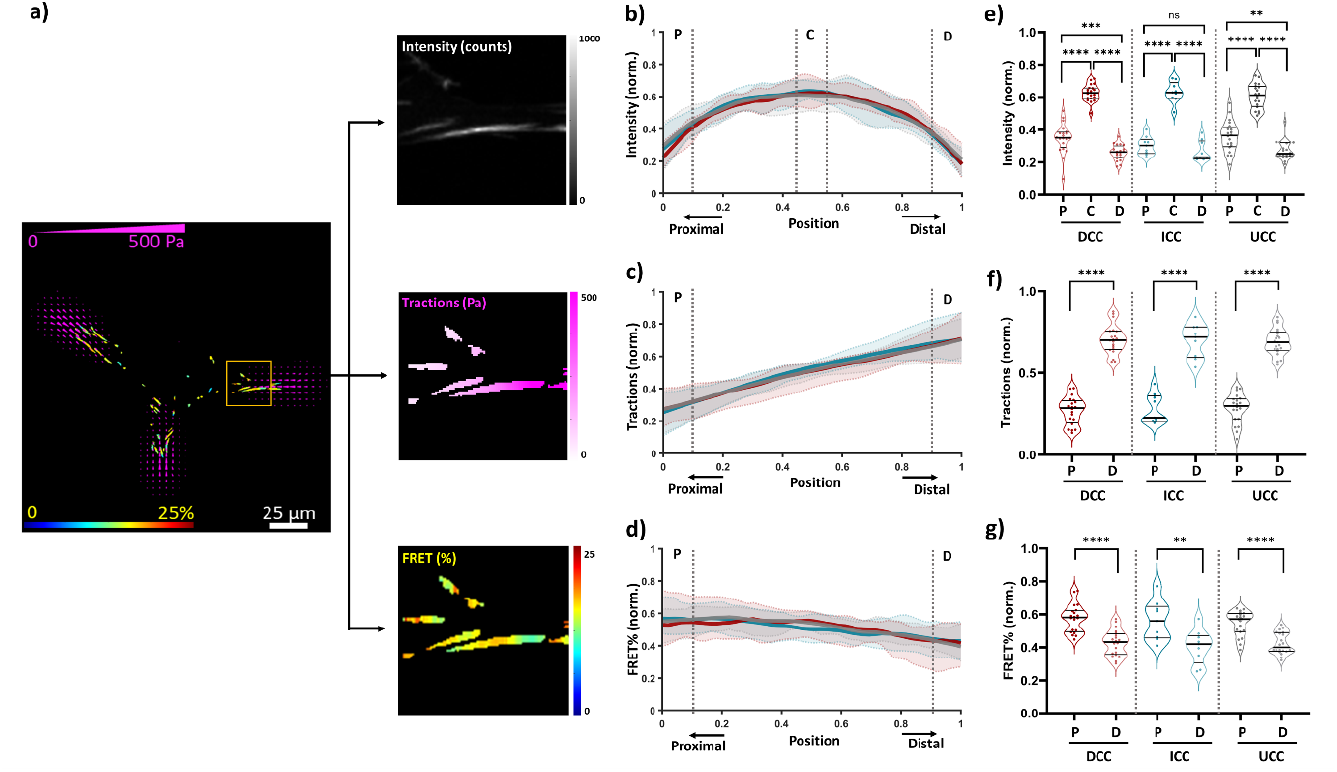
Spatial distribution of intensity, tractions, and FRET efficiency across focal adhesions. (**a**) Representative traction/FRET overlay image for sub-FA analysis; intensity (top), tractions (middle), and FRET efficiency (bottom) distribution along the length of individual FAs shown in the yellow box. (**b-d**) Line plots of the distribution profiles of normalized intensity (b), tractions (c), and FRET efficiency (d) along the proximal-to-distal position on FAs. Data of individual FAs (Supplementary Fig. 3) were normalized (between zero and one) and pooled per cell. Cell-averaged data were again split into three groups: directly correlated (DCC, blue), inversely correlated (ICC, red), and uncorrelated (UCC, gray). Shaded regions represent the distribution of cell-averaged data within the 10^th^ and 90^th^ percentiles; thick lines represent the global average of DCC, ICC and UCC. Proximal 10% (P), central 10% (C), and distal 10% (D) regions of FAs are delineated by dashed lines. (**e–g**) Statistical comparison of the proximal, central, and distal regions is shown for intensity (e), while only proximal and distal regions were compared for traction (f) and FRET efficiency (g). n_DCC_ = 20 cells, 630 FAs; n_ICC_ = 9 cells, 240 FAs; n_UCC_ = 22 cells, 638 FAs. Statistical analysis between two groups were performed using unpaired two-tailed parametric Student’s t-tests: *p < 0.05, **p < 0.01, p*** < 0.001 and ****p < 0.0001.

Despite differences in the correlation patterns at the FA level, the spatial profiles of vinculin density, tractions and vinculin tension within individual FAs exhibited a consistent polarity. Vinculin density peaked at the center of FAs, with fluorescence intensity significantly higher in this region compared to the proximal and distal ends (Fig. 5b and e). Tractions and vinculin tension were both significantly higher at the distal tip, the region farthest away from the cell center (Figs. 5c-g), suggesting a conserved relationship between these two variables at the sub-FA level. However, while tractions peaked sharply (Fig. 5c), FRET efficiency remained stable at the proximal end and gradually decreased towards the distal end, indicating increased vinculin tension mainly in the distal regions (Fig. 5d). Remarkedly, this spatial organization was observed across directly correlated, inversely correlated, and noncorrelated groups, highlighting the universal nature of this functional polarity. Sub-FA spatial analysis, performed separately for coupled and uncoupled FAs, showed similar patterns (Supplementary Fig. 3), suggesting that vinculin tension and traction distribution are conserved, independent of whether an FA actively contributes to significant traction generation.

## Discussion

This work introduces an integrated approach for investigating FA mechanics, combining simultaneous measurements of molecular tension and cellular tractions. By improving experimental sample preparation, streamlining data acquisition, and developing custom software, we achieved a highly detailed analysis of FA forces. These methodological advancements allowed us to explore the spatial coordination of cellular traction and vinculin tension, revealing new insights into the adaptability of FAs in force regulation and mechanotransduction.

Fluorescence imaging of cells expressing the VinTS enabled us to examine how FA morphology and vinculin content influence cellular tractions and molecular tension. Smaller FAs exhibited a broad spectrum of vinculin tension, ranging from very low to very high, while larger FAs were predominantly associated with moderate vinculin tension. This observation aligns with earlier findings that smaller FAs experience higher molecular tension, which decreases to average levels as FAs mature^25^. FAs with very low vinculin tension, on the other hand, were mainly located at the cell center, where FAs are typically smaller and subject to compression forces^28^. Interestingly, vinculin tension was inversely correlated with vinculin density, suggesting that higher vinculin packing reduces the tension per vinculin molecule. A weaker but similar trend was observed for total vinculin content, indicating that FAs may distribute larger forces across more vinculin molecules as the content increases. To support this idea, we found that larger FAs and those with higher vinculin content and density were associated with higher tractions. While similar correlations between tractions and FA properties such as size and intensity have been previously reported^47–49^, this correlation was only valid for nascent and growing FAs at the cell’s leading edge, not for trailing or disassembling FAs^50^. Moreover, according to Zhou et al., FA properties like area or vinculin content are insufficient to predict tractions and tension without considering the FA’s maturation history^51^. Our findings align with the conclusions of these studies: small FAs with low vinculin content exhibited significant variability in tractions and vinculin tension, reinforcing the importance of FA maturation and context. Unfortunately, due to the heterogeneous shape of the fibroblasts in this study, it was not possible to differentiate between leading and trailing FAs. Future studies using time-lapse imaging combined with TFM-FRET approach could provide clearer insights into how vinculin tension and tractions change throughout the FA lifecycle.

Considering the roles of actomyosin contractility and FA properties in regulating tractions^52–55^ and vinculin recruitment^13,17,56,57^, we investigated whether a relationship exists between vinculin tension and traction at the FA level. While no correlation between FA-averaged vinculin tension and traction was observed when cells were pooled, analyzing individual cells revealed significant cell-to-cell variability. Nearly half of the cells showed a direct correlation, where higher FA vinculin tension corresponded to higher tractions. A smaller subset, however, exhibited an inverse relationship, where decreased FA tension corresponded to elevated tractions. This variability likely reflects differences in the way cells interact with the ECM. We propose that highly engaged cells, such as those actively spreading or migrating, exhibit a direct correlation between FA tension and traction. In contrast, cells with deteriorating cell-ECM interactions, such as those retracting or shrinking, may display an inverse correlation. In these cases, tension is released from peripheral FAs while central FAs remain under tension. This observation aligns with prior studies indicating that peripheral FAs are more dynamic and prone to disassembly compared to central FAs^27^. Retracting FAs, which are linked to concentrated tractions before disassembly^8,50^, are known to experience lower vinculin tension^25^. Notably, a significant portion of cells exhibited no clear correlation between FA tension and traction. This may suggest that these cells were in a stable, non-migratory state, with with less dynamic cell-ECM interactions during imaging or that their FAs were in mixed functional phases, with some leading FAs promoting spreading and others trailing during retraction. These observations emphasize the complexity of FA dynamics, where both direct and inverse correlations emerge based on the state of the cell and FA behavior. Combining our TFM-FRET approach alongside tools to modulate acto-myosin activity, such as ROCK Inhibitor Y27632 or MLCP inhibitor Calyculin A, combined with time-lapse imaging, could provide deeper insights into these dynamic relationships.

Recognizing cell-to-cell variability in the relation between vinculin tension and traction, we investigated whether these two exhibit distinct correlations at sub-FA level. In agreement with earlier works^9,24,29,30^, sub-FA spatial analysis revealed distinct patterns of vinculin density, traction, and tension along the FA length, reflecting a mechanism for force transmission that is independent of cellular state. In the proximal half of the FAs, closer to the cell center, vinculin density gradually increased, driving an increase in traction despite the low and stable molecular tension. In contrast, the distal half, closer to the cell periphery, showed a decreasing vinculin density alongside an increasing vinculin tension, suggesting that higher vinculin tension compensates for reduced vinculin density to sustain force transmission. These results suggest a spatial coordination between vinculin density and tension to regulate tractions, highlighting the adaptability of FAs in force generation and mechanotransduction. Unexpectedly, unlike the FA-level results, vinculin density and tension were not anticorrelated at sub-FA level. Previous studies using the vinculin conformation sensor (VinCS), have shown that vinculin adopts an open, active conformation more frequently at the proximal end of FAs ^6,10^. This suggests that vinculin density alone may not regulate tension; instead, the amount of activated vinculin, capable of engaging in force transmission, could be the key factor. We hypothesize that the molecular tension is lower at the proximal end, where activated vinculin is more abundant, and higher at the distal end, where vinculin activation decreases. Overall, these results indicate that vinculin tension is regulated at two levels: vinculin density determines overall tension at the FA level, while vinculin activation controls tension distribution at the sub-FA level. To explore this further, future studies could combine TFM-FRET with VinCS or semi-digital vinculin tension probes^31,58,59^ to investigate how vinculin activation state and its spatial distribution shape the relationship between vinculin tension and tractions.

The substantial scatter in the data across all experiments in this study suggests that a linear relationship between mean traction and vinculin tension may not be feasible. Tractions likely depend on the combined effect of vinculin tension, density and activation, rather than vinculin tension alone. For instance, low vinculin tension paired with high active vinculin density could generate tractions similar to high tension with low active vinculin density. Additionally, vinculin tension and tractions may not be temporally synchronized. Vinculin tension fluctuates on short timescales^24^, while tractions, as large-scale outputs, take longer to adjust. Since stress-state bead images for TFM were acquired before FLIM imaging, vinculin tension changes during FLIM acquisition might not be reflected in the traction data. Finally, while vinculin plays a key role in FA stability and maturation, it is dispensable for FA formation and traction generation^13,60^. Exploring other FA proteins, such as talin or integrin, may yield stronger correlations with traction forces in future studies.

In conclusion, this study presents a novel approach for simultaneously measuring tractions and molecular tension within FAs, enabling new insights into their mechanics. We showed that the relationship between vinculin tension and traction depends on cellular states and FA context. Sub-FA analysis revealed a spatial coordination of vinculin density and tension, allowing FAs to balance forces and sustain effective traction. These findings highlight the adaptability of FAs in mechanotransduction and underscore the potential of this approach for uncovering the dynamic and spatial regulation of forces within FAs under diverse cellular conditions.

## Materials and Methods

### Cell Culture and Transient Transfection

Vinculin^-/-^ mouse embryonic fibroblasts (MEF^-/-^) were kindly provided by Wolfgang H. Goldmann and Ben Fabry^61^ and maintained in Dulbecco’s Modified Eagle Medium with 4.5 g/L D-glucose (Gibco) supplemented with 10% v/v fetal bovine serum (Sigma-Aldrich), 1% GlutaMax (Thermo Fisher), 0.1% Gentamycin (Carl Roth), 1% v/v Sodium Pyruvate (Sigma-Aldrich) and 1% v/v Non-Essential Aminoacids (Sigma-Aldrich) at 37°C, 20% CO2 and 5% oxygen. The cells were passaged at 80-90% confluency. Plasmid DNA expressing the vinculin tension probe VinTS-tCRMod-GGSGGS7 (Addgene #111764), and its control tCRMod-GGSGGS7 (Addgene #111761), were gifts from Brenton Hoffman^29^. Plasmid DNA expressing free Clover (Addgene #40259) was a gift from Michael Lin^62^. 24 hr after seeding on 29 mm dishes, cells were transfected using Mirus TransIT-X2 Dynamic Delivery System (Mirus Bio, MIR 6006) according to the manufacturer’s instructions. Per dish, 300 ng of plasmid DNA and 1 μL of TransIT-X2 were mixed in 100 μL of serum-free DMEM. The solution was incubated for 15 mins at RT and then delivered on the cells dropwise. Cells were left to grow overnight for expression of the Vinculin tension probes before being transferred to the collagen coated polyacrylamide hydrogels.

### Substrate Preparation

Polyacrylamide hydrogels of different stiffnesses were prepared according to previously reported protocols^39,63^. Briefly, all reagents were brought to room temperature and gels with a Young’s modulus of 4.5 kPa or 13 kPa were prepared by diluting 40% acrylamide (Sigma-Aldrich, A4058) down to 5% or 6.8% v/v and 2% bisacrylamide (Sigma-Aldrich, M1533) down to 0.15% or 0.17% v/v respectively in milliQ water. The mixture was supplemented with 1:60 dark-red fluorescent beads (Invitrogen, FluoSpheres™ Carboxylate-Modified Microspheres, 200 nm), vortexed and sonicated for 5 minutes. The polymerization of the gel was initiated by ammonium persulfate (4.4 µM) and catalyzed by N,N,N’,N’-tetramethylethylenediamine (0.1% v/v). The mixture was swiftly transferred to glass bottom dishes (Cellvis, D35-28-0-N) pre-treated with bind-silane and equipped with a 120 µm thick spacer (Invitrogen, Secure-Seal™, S24736), flattened with a siliconized cover glass and left to polymerize at room temperature for 45 minutes. After overnight incubation at 4°C in PBS, the gel surface was functionalized by two rounds of 0.4 µM Sulfo-SANPAH (Sigma-Aldrich, 803332) treatment under UV-irradiation with 365 nm wavelength emission for 15 minutes followed by a 2 hour incubation at 37°C in 20 µg/mL collagen type I (Gibco, A1048301) diluted in 0.1% v/v acetic acid at pH 3.4 for homogeneous coating^34^. ~30,000 transfected MEFs were seeded per dish and allowed to spread overnight before imaging.

### Mechanical Characterization of PAA gels

The rheological properties of the polyacrylamide hydrogels were acquired using a stress-controlled MCR702 rheometer (Anton Paar; SN 81385195) in combination with the RheoCompass software (Anton Paar; V1.30.1064). The solvent plate (Anton Paar, P-PTD200; SN 83604799) was attached to a Peltier device to maintain a temperature of 37 °C and customized with a homemade roughened aluminum surface. A disposable top plate (Anton Paar, PP08; SN000) with an 8 mm diameter rough aluminum insert was employed. Polyacrylamide hydrogels were prepared and functionalized ex-situ exactly as described above, except for the use of two siliconized glass surfaces with 3D printed isolators to facilitate the separation of the polymerized gel discs (7 mm diameter and 1 mm in height prior to swelling) and their transfer from the glass to the rheometer. Following placement of the gel on the bottom plate, the upper plate was lowered at a constant rate until contact was reached and a normal force of 0.05 N or 0.25 N was achieved, for 4.5 kPa and 13 kPa gels, respectively. A waiting step was used for adding 1X PBS buffer until the gel was submerged, to avoid dehydration during measurements. A time sweep was performed for 180 seconds at an oscillatory (engineering) shear strain amplitude of 1% (verified to be within the linear region) and angular frequency of 1 rad/s. For the frequency sweep the strain amplitude was kept at 1% for a varied frequency from 100-0.1 rad/s and for strain sweep tests the strain amplitude was varied from 0.01% to 100% at an angular frequency of 1 rad/s. For all conditions at least three independent gels were measured. The Young’s modulus ***E*** was calculated from the frequency-independent shear modulus ***G*′** with the following equation:

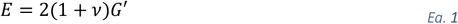

where we assumed a value of 0.49 for the Poisson’s ratio ***v***^39,64^.

### Fluorescence Microscopy Imaging

All live cell imaging for TFM and FRET measurements were carried out on a Leica TCS SP8 confocal fluorescence microscope equipped with FALCON (FAst Lifetime CONtrast) module. Prior to the imaging, samples were washed and submerged in warm live cell imaging solution (Invitrogen, A59688DJ). For TFM, confocal images were acquired using a 63X HC PL APO 1.2 NA water objective using variable zoom at a 1024 × 1024 resolution at 1AU. Excitation wavelengths were set to 488 nm (cell channel) and 633 nm (beads channel) using a pulsed laser at 20 MHz pulse frequency, 4 µW or 3 µW laser power respectively and 200 Hz unidirectional scanning speed. The detection ranges for the hybrid photomultiplier detectors were 495-600 nm and 640-800 nm respectively. Stressed state bead and cell images were collected as Z-stacks with 10 µm total thickness and 1 µm step size. All FRET images were acquired using the same objective but using a fixed 1.5X zoom at a 512 × 512 resolution (pixel size = 180 nm). The excitation wavelength was 488 nm (donor channel) and 561 nm (acceptor channel) using a pulsed laser at 20 MHz pulse frequency, 7 µW laser power and 50 Hz unidirectional scanning speed. For detection, a standard hybrid photon counting detector (Leica Microsystems GmbH) was set to 495-540 nm (donor channel) and 565-635 nm (acceptor channel) respectively. Leica FALCON module was used to process the photon arrival times and photon counts. Cell images were collected sequentially for the donor channel (17 frames) and for the acceptor channels (5 frames). After collecting all stressed state TFM and FRET images, one drop of 5% v/v SDS is placed on the sample and left for 15 minutes to remove the cells and allow the PAA hydrogel to return to a stress-free state. The relaxed state bead images were acquired using identical settings as for the stressed state bead images.

### 2D TFM Analysis

To calculate displacement and traction fields, TFM images were analyzed using in-house developed Matlab code based on previously published code^40^. Briefly, far-red bead channel images were filtered using a Gaussian filter to suppress noise and cell masks were acquired by thresholding. Spatial shifts between stressed and relaxed TFM images were corrected by means of rigid image registration with function *findshift* from the DipImage library^65^. To obtain the cell-induced movements of the beads, a free-form deformation image registration was applied to the bead channel^40^ using the open-source Elastix toolbox^66^. Tractions were recovered by considering PAA gels as homogeneous, isotropic, linear elastic half spaces. Then a modified Tikhonov regularized Fourier transform traction cytometry algorithm was applied, to avoid unwanted effects of the Young’s modulus of the substrate in the regularization parameter^39^. Since noise levels in the measured displacement fields were relatively similar throughout the dataset, we fixed a value for the regularization parameter that reduced noisy traction maps with limited smoothing. This value was used for all of the dataset. Mean tractions in Fig. 2c were calculated using the following formula:

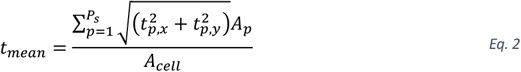

where ***P***_***s***_ denotes the number of pixels that constitute the traction footprint of the cell, ***t***_***p***,***x***_ and ***t***_***p***,***y***_ are the x and y components of traction vector at pixel ***p***, respectively, ***A***_***p***_ is the pixel area (~22.5 x 10^-3^ μm^2^), ***A***_***cell***_ is the area of the cell mask.

## FLIM-FRET Analysis

For the FRET analysis, fluorescence decays of each pixel is transformed into phasor space by the integrated Phasor module of SP8 FALCON. Shortly, the full decay ***I*(*t*)** is Fourier transformed into a real part (***G***) and an imaginary part (***S***) using:

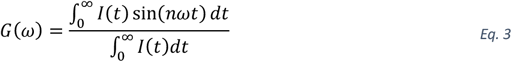

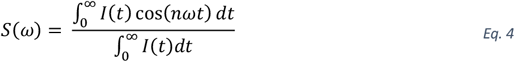

where ***n*** is the chosen harmonic (in this case, 1), ***ω*** is **2*πf*** and ***f*** is the repetition rate of the pulsed laser (in this case, 20 MHz). ***G*** and ***S*** values corresponds to the coordinates on the phasor space per pixel, effectively creating a 2D-histogram representation of all the pixel lifetime information of the image. Resulting ***G*** and ***S*** coordinates were exported for further analysis using a custom matlab script for phasor-based FRET calculations (Fig. 1c). The position of Clover (fluorescence lifetime ~3.0 ns, averaged from 11 MEFs expressing cytosolic Clover) was used to define the starting position of the FRET efficiency ttrajectory line. An approximate position of the background signal (***G* = 0.5, *S* = 0.5**), as observed from mock-transfected cells, was used to define the end point of the trajectory. Finally, each point on the phasor space, corresponding to a pixel on the image, was assigned the FRET value of the closest point on the FRET trajectory line, resulting in the FRET image. To remove the noise in the data, and to smooth the FRET efficiency distribution, a median convolution filter of 9 × 9 pixels was applied to the FRET image, as described previously by Ranjit et al. (2018)^43^.

## FA structural and molecular properties

For FA segmentation, binary masks were generated using Ilastik machine learning segmentation tool^46^ using the acceptor (mRuby2) channel images. Binary mask, FRET image and acceptor (mRuby2) intensity data were imported into MATLAB and used to compute mean FRET index, total and mean intensity per FA. Total FA intensity, given as counts, was used as indicator of total vinculin content, while average FA intensity, given as counts per µm^2^, was used as indicator of vinculin density; i.e. amount of vinculin per unit area. The function *bwconncomp* was used to obtain FA area and eccentricity. Before extracting TFM metrics, function *imresize* with bicubic interpolation was used to resample the TFM data to match the pixel size of the FLIM data. Then, rigid image registration of the TFM cell channel and FLIM donor channel was used to calculate the relative shift needed to spatially align both datasets. Lastly, all images were zero padded and/or cropped to bring all datasets to the same logical size. Coupled and uncoupled FAs were segregated by thresholding the displacement field and evaluating if they were part of the active regions (Fig. 4c). Mean traction per FA was calculated similar to Eq. 2:

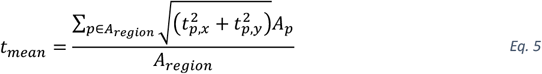

where ***A***_***region***_ is the area of a FA.

We developed the following algorithm to analyze acceptor intensity, FRET index and traction profile along FA longitudinal axes (Fig. 5, Supplementary Fig. 4). First, function *regionprops* (with input property *orientation*) was applied on each FA to obtain its principal direction, ***d***. A line, ***L***, passing through the FA centroid with direction *d* was drawn to obtain its two intersection points with the FA boundaries. The point closer to the cell mask centroid was selected as the initial point for the analysis. A pixel-by-pixel iterative process sampled and averaged the data (acceptor intensity, FRET index or tractions) within the FA mask along perpendicular lines ***p*** to ***L*** using an analogous formula to Eq. 2. The resulting data vector was linearly interpolated using function *interp1* into a normalized 50 point axis (see Fig. 5**Figure *5***). FAs with a length smaller than 9 pixels were discarded to avoid sharp discontinuities. To obtain the FA orientation with respect to the cell, we defined the direction ***r*** as the line that connected the centroid of the cell with that of the FA and calculated its acute angle with ***d*** (Fig. 3e,i).

## Statistics and Reproducibility

At least three independent technical replicates were performed for each experiment, and biologically independent samples were analyzed. The experiments were not randomized and researchers were not blinded during experiments and outcome assessment. Data quantification was conducted using automated tools. Graphs were generated using MATLAB R2022b or GraphPad Prism 10.4.1 (GraphPad Software). Sampled cells were excluded only if they did not meet the requirements for traction recovery (e.g. if the displacement field was unreliable due to large stage shifts between the stressed and relaxed state or if the assumption of a displacement/stress free boundary was violated). Normality of the data was assessed using the Shapiro-Wilk test. For data following a normal distribution, a two-tailed, unpaired, parametric Student’s t-test was applied, while the nonparametric Mann-Whitney U-test was used for non-normally distributed data. Statistical significance was defined as follows: *p< 0.05, **p < 0.01, ***p < 0.001, ****p < 0.0001. Simple linear regression was used to assess relationships between variables. The goodness-of-fit (R^2^) was calculated to evaluate the strength of the relationships. Further statistical details for individual experiments are provided in the corresponding figure legends.

## Code availability

All custom written codes will be made available at a Zenodo repository.

## Data Availability

All data, including the raw microscopy files and the source data for charts and graphs, is available from the corresponding authors upon request.

## Acknowledgments

The authors are grateful to the following funding sources: S.R and S.A. were supported by KU Leuven internal funding: IDN/20/021, KA/20/026 and C14/22/085. S.D. was supported by KU Leuven internal funding: ZB/22/028. S.R. and H.V.O. received funding from the Research Foundation Flanders (FWO) through a project with grant number G0C2422N. H.V.O. and L.K were supported by iBOF project 21/083 C. H.V.O. received the FWO infrastructure grant I009718N. In addition, S.A. and J.B.F. are recipients of FWO fellowships with grant numbers 1S95125N and 1259223N respectively.

## Author contributions

Experiments and data analysis were performed by S.A and L.K. The rheological characterization of the hydrogels was performed by D.L. and S.D., under the supervision of R.C. and H.V.O. The customized Matlab algorithms were developed by L.K., Q.C. and J.B.F.. Q.C., M.C., H.V.O. and S.R. developed the initial concept. The work was supervised by H.V.O. and S.R. The original draft was written by S.A., under the supervision of S.R. All authors contributed to the writing of the final version.

## Supplementary Figures

**Supplementary Figure 1:**
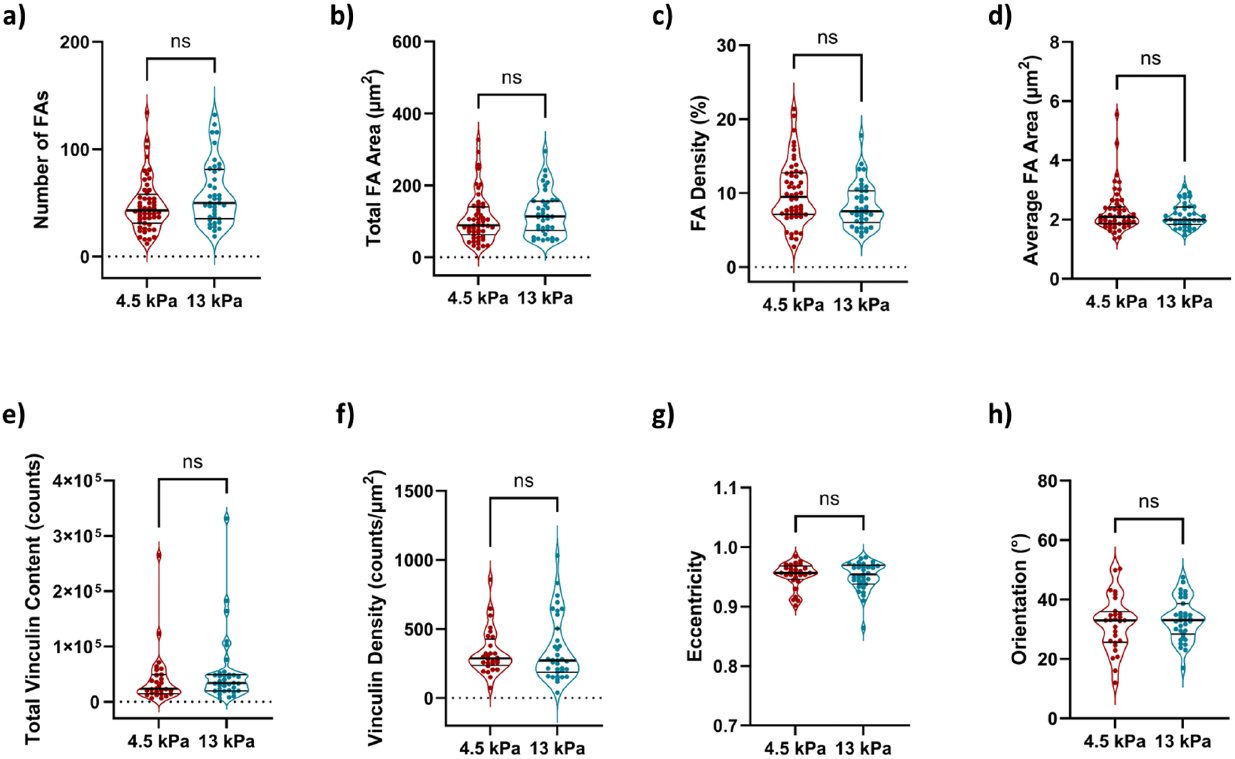
Effect of the stiffness on FA structural and molecular properties. Total number of FAs (**a**), total surface area of FAs (**b**), FA density (i.e. percentage of the cell occupied by FAs) (**c**), cell-averaged FA area (**d**), total vinculin content (i.e. total acceptor channel intensity) (e), average vinculin density (i.e. acceptor channel intensity per unit area) (f), cell-averaged eccentricity (**g**) and FA orientation relative to the cell center (**h**) were compared between soft (4.5 kPa) and stiff (13 kPa) substrates. n = 52 (4.5 kPa) and 40 (13 kPa) for (a-d). n = 28 (4.5 kPa) and 35 (13 kPa) for (e-h). Statistical analysis between groups was carried out using nonparametric Mann-Whitney U-test.

**Supplementary Figure 2:**
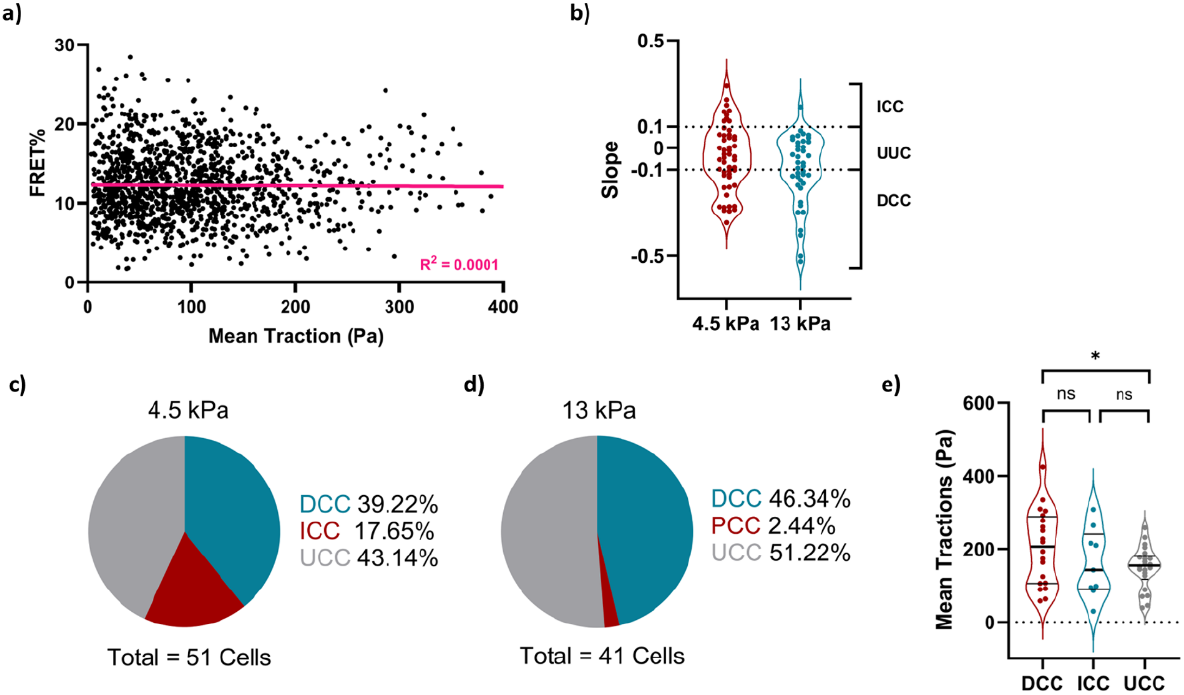
Cell-level comparison of mean FA traction and mean FRET efficiency. (**a**) Mean traction of FAs were compared to mean FRET efficiency of FAs when all cells on soft substrate (4.5 kPa) included in the analysis. n = 1508 FAs from 51 cells. Purple line represents the linear regression fit, with R^2^ value indicated on the graph. (**b**) Distribution of linear correlation slopes for soft (4.5 kPa) and stiff (13 kPa) substrate, obtained from FRET vs Mean Traction plots of each cell after the maximum FRET and maximum traction values were normalized to 1. n = 51 (for 4.5 kPa) and 40 (for 13 kPa) cells. (**c-d**) Percentile distribution of directly correlated (DCC; slope below −0.1), inversely correlated (ICC; slope above +0.1) and uncorrelated (UCC; slope between-0.1 and +0.1) cells for soft and stiff substrates. (**e**) Cell-averaged tractions of DCC, ICC and UCC compared for soft substrate. n = 20, 9 and 22 cells, respectively. Statistical analysis between groups was carried out using parametric, unpaired, two-tailed Student’s t-test. *p < 0.05.

**Supplementary Figure 3:**
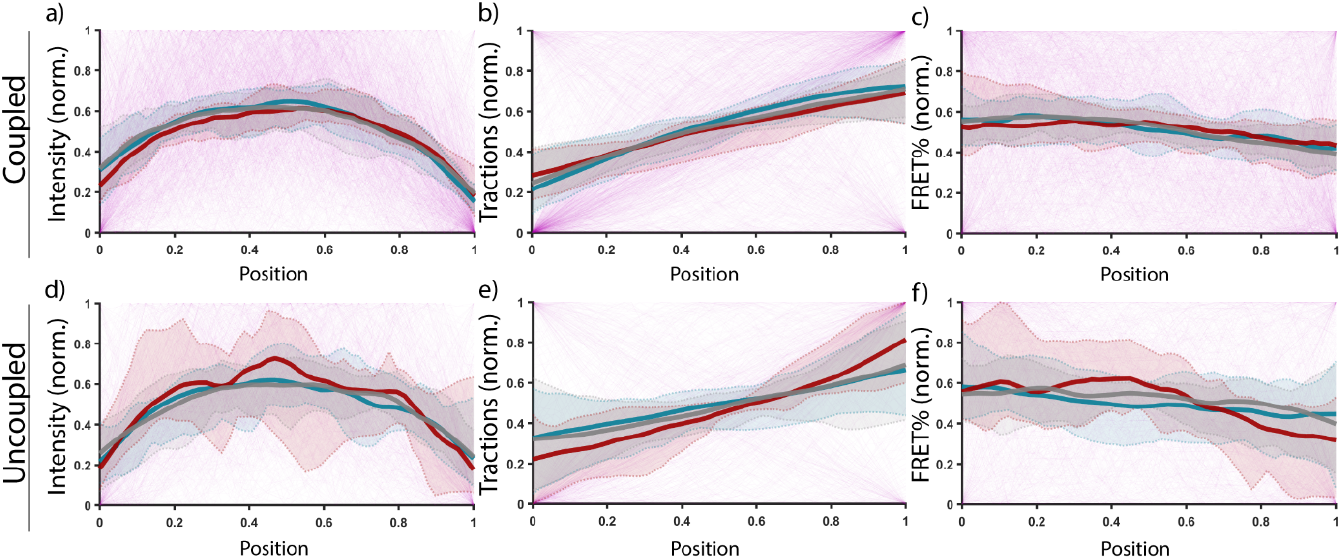
Intensity, traction and FRET distribution of coupled and uncoupled FAs. Sub-FA spatial distribution of intensity, traction and FRET, compared between directly correlated cells (DCC; blue), inversely correlated cells (ICC; red), and uncorrelated cells (UCC; gray), as well as between the coupled (**a-c**) and uncoupled FAs (**d-f**). Distributions for individual FAs are represented as thin lines (DCC + ICC + UCC; purple). n_DCC_ = 20 cells, 406 coupled and 224 uncoupled FAs; n_ICC_ = 9 cells, 203 coupled, 37 uncoupled FAs; note the small sample size of ICC uncoupled FAs. n_UCC_ = 22 cells, 406 coupled and 232 uncoupled FAs.

**Supplementary Figure 4:**
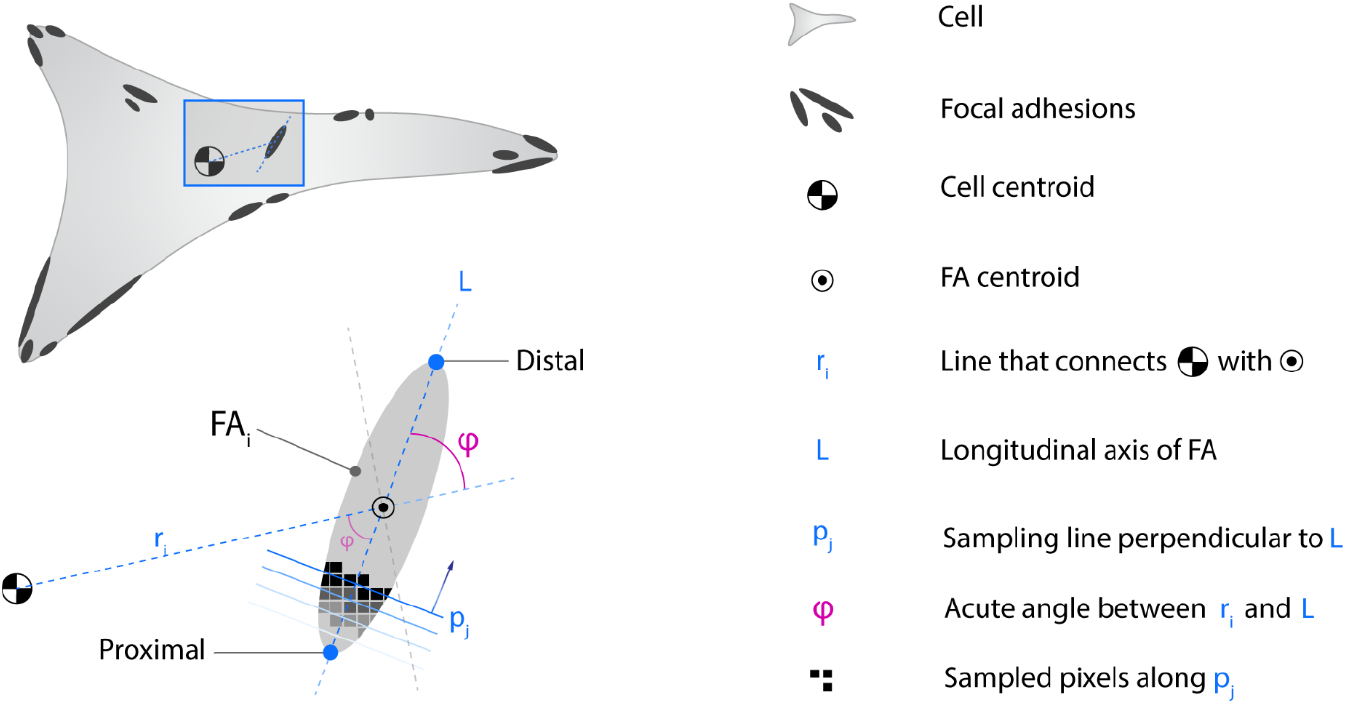
Sampling approach to determine the spatial distribution of sub-FA metrics. For every i^th^ FA, we defined a line ***L*** oriented in the principal direction of the FA and crossing the FA’s centroid. The proximity of the endpoints of L to the cell’s centroid determined if an endpoint is considered proximal (closest) or distal (farthest). The line that connects the cell’s centroid to the centroid of the i^th^ FA is defined as ***r***_***i***_. The acute angle (< 90°) between ***r***_***i***_ and ***L*** was defined as ***φ***. The sub-FA spatial distribution of intensity, traction and FRET were acquired using a pixel-by-pixel iterative approach. Starting at the proximal end of ***L***, a perpendicular line ***p***_***j***_ to ***L*** was created and all pixels coinciding with this line were sampled, the values averaged and stored. Next, the line ***p***_***j***_ was moved one pixel closer towards the distal end along ***L*** to create a new line ***p***_***j*+1**_ and the sampling process was repeated until line ***p*** coincided with the distal endpoint of ***L***.

