## Supporting Information for "Linking Molecular Tension and Cellular Tractions: A Multiscale Approach to Focal Adhesion Mechanics"

### Supplementary Figures

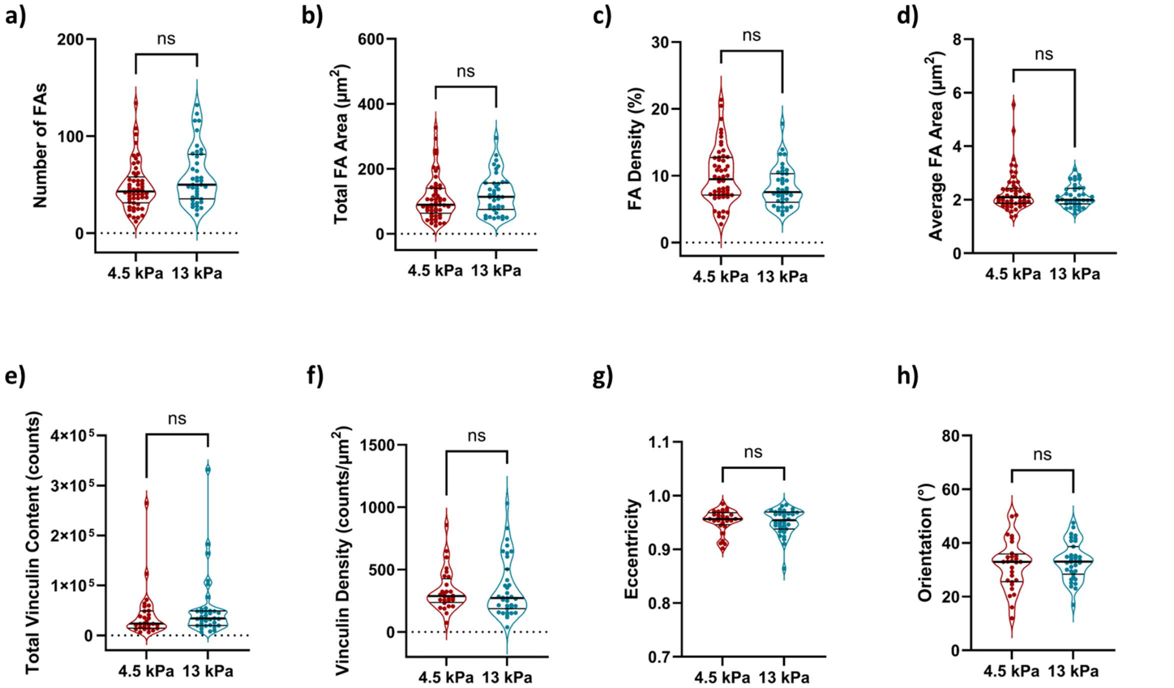


**Supplementary Figure 1: Effect of the stiffness on FA structural and molecular properties.** Total number of FAs (**a**), total surface area of FAs (**b**), FA density (i.e. percentage of the cell occupied by FAs) (**c**), cell-averaged FA area (**d**), total vinculin content (i.e. total acceptor channel intensity) (**e**), average vinculin density (i.e. acceptor channel intensity per unit area) (**f**), cell-averaged eccentricity (**g**) and FA orientation relative to the cell center (**h**) were compared between soft (4.5 kPa) and stiff (13 kPa) substrates. n = 52 (4.5 kPa) and 40 (13 kPa) for (a-d). n = 28 (4.5 kPa) and 35 (13 kPa) for (e-h). Statistical analysis between groups was carried out using nonparametric Mann-Whitney U-test.

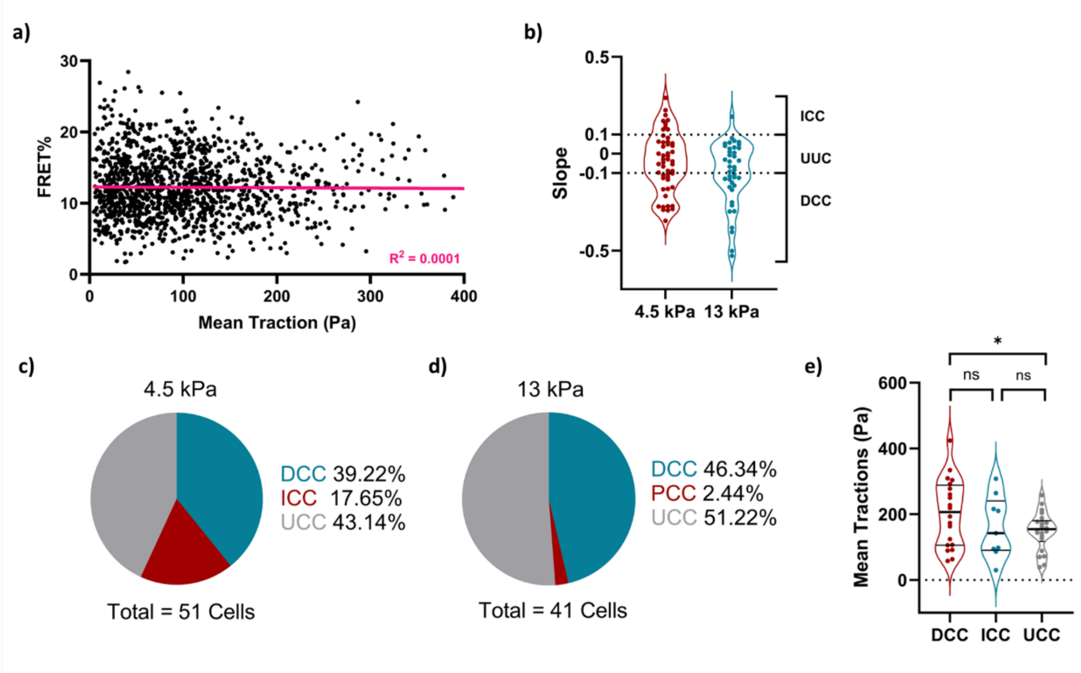


**Supplementary Figure 2: Cell-level comparison of mean FA traction and mean FRET efficiency.** (**a**) Mean traction of FAs were compared to mean FRET efficiency of FAs when all cells on soft substrate (4.5 kPa) included in the analysis. n = 1508 FAs from 51 cells. Purple line represents the linear regression fit, with $R^2$ value indicated on the graph. (**b**) Distribution of linear correlation slopes for soft (4.5 kPa) and stiff (13 kPa) substrate, obtained from FRET vs Mean Traction plots of each cell after the maximum FRET and maximum traction values were normalized to 1. n = 51 (for 4.5 kPa) and 40 (for 13 kPa) cells. (**c-d**) Percentile distribution of directly correlated (DCC; slope below -0.1), inversely correlated (ICC; slope above +0.1) and uncorrelated (UCC; slope between -0.1 and +0.1) cells for soft and stiff substrates. (**e**) Cell-averaged tractions of DCC, ICC and UCC compared for soft substrate. n = 20, 9 and 22 cells, respectively. Statistical analysis between groups was carried out using parametric, unpaired, two-tailed Student's t-test. $*p < 0.05$.

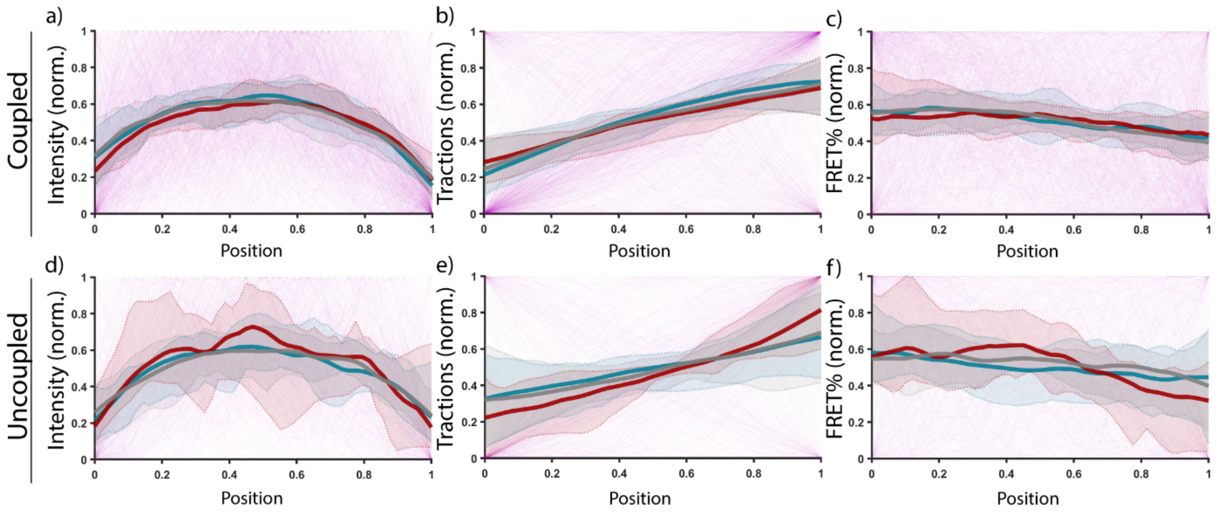


**Supplementary Figure 3: Intensity, traction and FRET distribution of coupled and uncoupled FAs.** Sub-FA spatial distribution of intensity, traction and FRET, compared between directly correlated cells (DCC; blue), inversely correlated cells (ICC; red), and uncorrelated cells (UCC; gray), as well as between the coupled (**a-c**) and uncoupled FAs (**d-f**). Distributions for individual FAs are represented as thin lines (DCC + ICC + UCC; purple). $n_{DCC}$ = 20 cells, 406 coupled and 224 uncoupled FAs; $n_{ICC}$ = 9 cells, 203 coupled, 37 uncoupled FAs; note the small sample size of ICC uncoupled FAs. $n_{UCC}$ = 22 cells, 406 coupled and 232 uncoupled FAs.

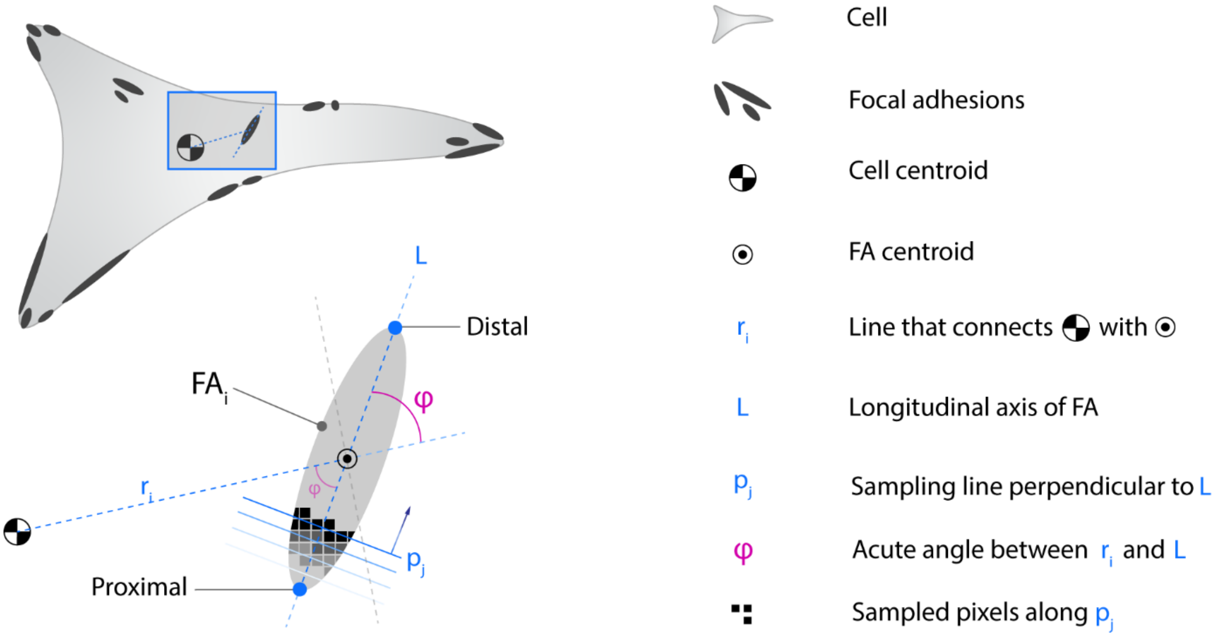


**Supplementary Figure 4: Sampling approach to determine the spatial distribution of sub-FA metrics.** For every i$^{th}$ FA, we defined a line $L$ oriented in the principal direction of the FA and crossing the FA's centroid. The proximity of the endpoints of L to the cell's centroid determined if an endpoint is considered proximal (closest) or distal (farthest). The line that connects the cell's centroid to the centroid of the i$^{th}$ FA is defined as $r_i$. The acute angle (< 90°) between $r_i$ and $L$ was defined as $\varphi$. The sub-FA spatial distribution of intensity, traction and FRET were acquired using a pixel-by-pixel iterative approach. Starting at the proximal end of $L$, a perpendicular line $p_j$ to $L$ was created and all pixels coinciding with this line were sampled, the values averaged and stored. Next, the line $p_j$ was moved one pixel closer towards the distal end along $L$ to create a new line $p_{j+1}$ and the sampling process was repeated until line $p$ coincided with the distal endpoint of $L$.
